# Detection of idiosyncratic gaze fingerprint signatures in humans

**DOI:** 10.1101/2023.09.18.558217

**Authors:** Sarah K. Crockford, Eleonora Satta, Ines Severino, Donatella Fiacchino, Andrea Vitale, Natasha Bertelsen, Elena Maria Busuoli, Veronica Mandelli, Michael V. Lombardo

## Abstract

Do individuals possess a ‘*gaze fingerprint*’ that reveals how they uniquely look at the world? We tested this question by examining intra- and inter-subject gaze similarity across 700 static pictures of complex natural scenes. Independent discovery (n=105) and replication (n=46) datasets revealed that gaze fingerprinting is possible at relatively high rates (e.g., 52- 63%) compared to chance (e.g., 1-2%). We also introduce the idea of a ‘*gaze fingerprint barcode*’, which can reveal how an individual can be gaze fingerprinted across a large array of stimuli. Pre-registered longitudinal follow-up experiments show that gaze fingerprint barcodes are stable within-individual over short (e.g., weeks to months) and long (e.g., years) time frames. Finally, we find that increased gaze fingerprintability is associated with decreased levels of autistic traits. Overall, this work showcases the potential of gaze fingerprinting and may help reveal precision biomarkers relevant for studying conditions with atypical gaze patterns, such as autism.

---

Human gaze patterns provide one of the most important signals regarding what an individual finds most salient about their environment. Many factors outside of the individual have been theorized to explain variance in gaze patterns, such as context, task, novelty, and characteristics of the stimuli themselves (Borji et al., 2013; de Haas et al., 2019; Einhauser et al., 2008; Guy et al., 2019; Henderson & Hayes, 2017; Itti et al., 1998; Xu et al., 2014). However, because gaze patterns can also be quite stable and systematic (Avni et al., 2020; Broda & de Haas, 2022; de Haas et al., 2019; Guy et al., 2019; Keles et al., 2022; Linka et al., 2022; Linka & de Haas, 2020), this suggests that there must also be a strong within-individual factor that explains variance in gaze patterns. Recent twin studies have revealed heritable common genetic variation to be one such powerful within-individual mechanism constraining gaze patterns (Constantino et al., 2017; Kennedy et al., 2017; Portugal et al., 2023). These insights suggest the possibility that gaze patterns may be an important individuating signature or ‘fingerprint’ of high neurobiological and developmental significance. With the advent and uptake of technology that could routinely collect eye gaze data at-scale in very large quantities, the ability to identify individuals by what they look at merged with other personal data could have very large impact and raise issues regarding privacy (Cantoni et al., 2018; Kröger et al., 2020; Liebling & Preibusch, 2014; Rigas et al., 2016). Gaze fingerprint markers may also be of high translational relevance for precision medicine approaches applied to conditions where gaze is atypical, such as autism. Atypical gaze patterns are evident in autism both in early development (Constantino et al., 2017; Jones & Klin, 2013) and at older ages (Wang et al., 2015). Such atypical social visual engagement in autism is theorized to be a manifestation of genetically constrained, yet atypical, biological niche construction (Johnson, 2017; Johnson et al., 2015; Klin et al., 2020; Lombardo et al., 2019; Shultz et al., 2018). Gaze fingerprinting markers could provide an individualized metric for underlying differential biology (Lombardo et al., 2019) and for assessing change as a function of manipulations to an individual’s environmental niche (e.g., early intervention).

In this study we investigate the phenomena of ‘gaze fingerprinting’ – that is, identification of an individual based on repeatable spatial distributions of gaze patterns that are unique to the individual. Prior work focusing on saccade-based metrics (e.g., velocity, acceleration, vigor) has shown some promise as an individuating biometric (e.g., (Bargary et al., 2017; Choi et al., 2014; Juhola et al., 2013; Rigas et al., 2016)). In contrast, utilization of gaze measures that better define where and what someone samples in their environment (i.e. spatial gaze distribution patterns) have been much less studied. Prior work examining spatial fixation density and/or ‘gaze heatmaps’ has shown that individualized spatial gaze patterns can be quite stable and systematic (Avni et al., 2020; Broda & de Haas, 2022; de Haas et al., 2019; Guy et al., 2019; Linka et al., 2022; Linka & de Haas, 2020). However, such work does not address whether the ‘target’ individual can be accurately identified when compared to gaze patterns of many other ‘distractor’ individuals.

To our knowledge, three prior studies have directly attempted to address whether individuals can be successfully identified solely based on spatial gaze patterns that define what they sample in complex visual environments (Keles et al., 2022; Kennedy et al., 2017; Rigas & Komogortsev, 2014). These studies suggest that some, though not all, individuals may be gaze fingerprintable and that such gaze fingerprints may be potentially driven in part by genetic similarity. For example, two independent studies examining a small number of dynamic movie stimuli have reported identification rates around 30-40% (Keles et al., 2022; Rigas & Komogortsev, 2014). Both studies noted limitations in using a small number of dynamic movie stimuli that do not allow for sampling gaze patterns across a large variety of possible stimulus categories and/or contexts. Furthermore, both studies also report observations that increasing the amount of gaze data sampled can lead to an increase in identification accuracy. These key limitations may suggest that sampling from a larger array of visual stimuli within-individual could lead to marked increases in our ability to gaze fingerprint an individual.

In contrast to movie stimuli, to our knowledge, no studies thus far have directly examined gaze fingerprinting the same individual on static pictures of complex scenes. Unlike with longer-duration movie viewing, a broad array of different categories and contexts can be sampled in a short period of time with rapid presentations of many static pictures of complex visual scenes (e.g., (de Haas et al., 2019; Xu et al., 2014)). A closely-related study to the concept of gaze fingerprinting showed that twin pairs could be accurately identified at a rate of 29% from a gaze patterns to small set of static pictures of complex scenes (Kennedy et al., 2017). In contrast, identification accuracy of dizygotic twin pairs was substantially lower 7%, but higher than non-twin pairs and chance accuracy of 1% (Kennedy et al., 2017). Given the near 30% accuracy rate for identifying twin pairs, it is possible that accuracy rates for static stimuli on the same individuals may be much higher than ∼30%, particularly when sampled in a big data context with a stimulus-rich experimental design. Because this study found a dissociation in identification rate between MZ and DZ twins, it is conceptually related to ‘gaze fingerprinting’ by demonstrating that the degree gaze similarity scales with genetic similarity. Furthermore, this twin study suggests that gaze similarity and gaze fingerprinting itself may be an eye tracking biomarker of something biologically more insightful regarding how heritable polygenic genomic architectures build early neural circuitry in ways that prepare it to seek out, explore, and construct similar environmental niches to develop within.

The current set of studies we have conducted considerably expands our understanding of the concept of gaze fingerprinting when expanded at-scale in a big data context such as stimulus-rich experimental design. Here we show a proof-of-concept that when static picture stimuli are sampled in a stimulus-rich fashion (e.g., 700 stimuli) across a wide range of semantic categories, that gaze fingerprinting shows much promise as an individualized marker of high biological significance, which may also be translationally relevant to conditions such as autism. We measured spatial distributions of gaze patterns with gaze heatmaps in a relatively large sample of participants across discovery and replication sets (discovery n=105; replication n = 46) viewing a large richly-annotated stimulus set of 700 complex scenes (Xu et al., 2014), repeated in two sessions separated by about 1-2 weeks. The unique ‘stimulus-rich’ feature of this study allows for dense sampling of gaze patterns across a large range of complex scenes depicting a variety of semantic features that have been heavily studied regarding visual salience (de Haas et al., 2019; Wang et al., 2015; Xu et al., 2014). This dense sampling of gaze across such a feature-rich stimulus set allows for gaze fingerprinting based on stimuli-averaged similarity, but also at-scale per each individual stimulus. Gaze fingerprinting at-scale on individual stimuli also allows for a novel concept we dub gaze fingerprint ‘barcoding’, whereby individuals are uniquely identifiable by their patterns of fingerprintable stimuli sets and the semantic features describing those sets. In a second set of pre-registered follow-up experiments, we tested whether gaze fingerprint barcodes are non-random within-individuals after a short-term (e.g., 19-94 days) or long-term (e.g., 1.87-5.16 years) delay. Finally, because gaze similarity is linked to genetically sensitive phenotypes such as autism (Avni et al., 2020; Constantino et al., 2017; Keles et al., 2022; Kennedy et al., 2017; Wang et al., 2015), we assessed whether gaze fingerprinting metrics are related to phenotypes underpinned by heritable common genetic mechanisms, such as autistic traits. We show that increasing level of autistic traits is related to less gaze fingerprintability. This result may be important under viewpoints about autism with respect to enhanced neuronal and behavioral noise that lead to idiosyncratic neural and behavioral phenotypes (Avni et al., 2020; Benkarim et al., 2021; Bleimeister et al., 2024; Bolton et al., 2018; Byrge et al., 2015; Cavallo et al., 2018; Dinstein et al., 2012, 2015; Hahamy et al., 2015; Hasson et al., 2009; Keles et al., 2022; Milne, 2011; Nunes et al., 2019; Pegado et al., 2020; Trakoshis et al., 2020).

## Materials and Methods

### Discovery dataset

The discovery dataset (labeled henceforth as ‘IIT’) comprised individuals tested in an eye tracking study carried out within the IIT Laboratory of Autism and Neurodevelopmental Disorders (IIT-LAND). This study was approved by the local ethics committee (Comitato Etico per le sperimentazioni cliniche dell’Azienda Provinciale per Servizi Sanitari della Provincia Autonoma di Trento; APSS). All participants gave written informed consent and were reimbursed for their time participating in the study. Participants were primarily recruited through social media and convenience sampling within the University of Trento Center for Mind and Brain Sciences (CIMeC) department. A total of n=120 typically-developing adult participants (age range 18-50 years) with normal or corrected-to-normal vision took part in this study. All participants took part in two sessions, whereby the separation between the two sessions was on-average 7 days (SD = 1.46 days). Due to missing gaze data for some individuals on some stimuli (e.g., participant looked away from screen), n=13 individuals were excluded from analysis if they had more than 15% of the 700 stimuli missing gaze data for either session 1 or session 2. A further n=2 participants were excluded due to delays between session 1 and session 2 that were over 20 days. The final sample utilized in all further analyses was n=105 (n=52 female, n=53 male). The sample had a mean age of 26.21 years (SD = 6.30) and was within the normal range on full-scale IQ (mean = 108.65, SD = 13.09, range = 75-138). While the main analyses on the IIT dataset use all individuals, in follow-up analyses (see Supplementary Figure 1) we split the IIT discovery dataset by sex in order to evaluate how fingerprint accuracy varies in smaller sample sizes (n=52 female, n=53 male) that are on-par with the size of the replication dataset (e.g., n=46).

### Replication dataset

As a replication dataset we re-analyzed anonymized publicly available test-retest eye tracking data reported from de Haas and colleagues (de Haas et al., 2019) (https://osf.io/n5v7t/). Henceforth, this replication dataset is referred with the original acronym given by de Haas and colleagues of ‘Gi’. The experimental design and stimuli for both the IIT discovery and Gi replication datasets were identical in that both utilized the OSIE dataset of 700 stimuli and participants were instructed to freely view the stimuli over a 3 second period in both test and retest sessions. The primary difference between the IIT discovery and Gi replication datasets was the duration of time in between test and retest sessions. While the IIT dataset had a delay of 7 days on-average (SD = 1.46), the delay in the Gi replication dataset a little more than double this duration (mean = 16 days, SD = 7 days). The sample size of the Gi dataset was n=48. However, n=2 individuals were excluded from analysis if they had more than 15% of the 700 stimuli missing gaze data for either session 1 or session 2. The sample characteristics of the Gi replication dataset indicate that the sample size was smaller (n=46) than the IIT discovery dataset (n=105). Furthermore, the Gi dataset was primarily female (e.g., n=11 male), whereas the sexes were relatively balanced in the IIT dataset.

### Eye Tracking Task and Stimuli

Participants were asked to freely watch 700 stimuli of complex natural scenes taken from the Object and Semantic Images and Eye-tracking (OSIE) stimulus set (Xu et al., 2014). Stimuli in the OSIE set are annotated both for their low pixel- and object-level features as well as higher-level semantic features (Xu et al., 2014). Semantic features represented categories such as face, emotion, touched, gazed, motion, sound, smell, taste, touch, text, watchability, and operability. Further annotation beyond these twelve categories was done to label stimuli by presence or absence of ‘humans’ or ‘animals’ as well as a categorization of whether the stimuli had ‘social’ (i.e. presence of human faces) or ‘non-social’ (i.e. absence of humans or animals) content. While all semantic features can be defined and analyzed independently, they co-occur within the OSIE stimuli in a non-independent manner. Therefore, to observe the emergent structure of co-occurrence of multiple semantic features as clusters, we utilized two-way hierarchical clustering with Jaccard distance matrices and ward.D2 as the clustering agglomeration method to reveal data-driven clusters across all 700 stimuli and across all semantic features (Figure 1A). This analysis reveals that there are 2 primary clusters amongst the 700 stimuli that largely define ‘social’ versus ‘nonsocial’ stimuli (Figure 1A). Semantic features can be separated into 6 clusters (Figure 1A).

**Figure 1:**
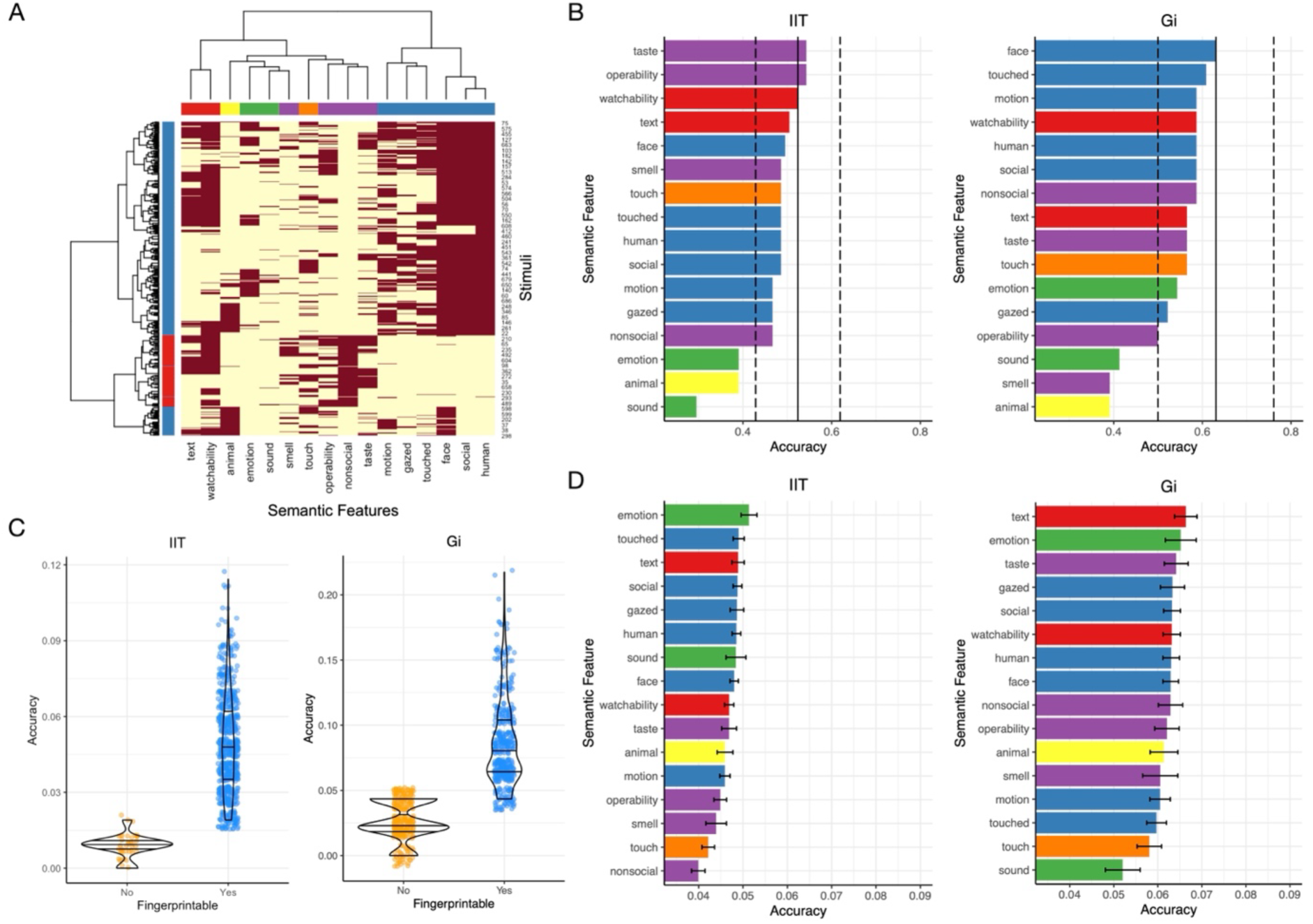
Gaze fingerprint accuracy on averaged gaze similarity across stimuli or for each individual stimulus. Panel A is a heatmap representation of the co-occurrence of various semantic features (columns) across all 700 stimuli (rows) in the OSIE stimulus set. Dark red indicates presence of a stimulus in a particular semantic feature category, while light yellow indicates absence of a stimulus in a particular semantic feature category. Colors for clusters of semantic features (underneath the column dendrogram) are used in panels B and D. The vertical black line in panel B shows gaze fingerprint accuracy (x-axis) when average gaze similarity is computed across all 700 stimuli (dotted vertical black lines represent bootstrap 95% confidence intervals). Each bar shows gaze fingerprint accuracy when computed as average gaze similarity on subsets of stimuli grouped by semantic features shown on the y-axis. On the left are accuracy values in the IIT discovery set, while on the right are the accuracy values for the Gi replication set. Panel C shows accuracy values (y-axis) for each of the 700 stimuli. The x-axis denotes stimuli that are fingerprintable or not. This distinction is made based on whether the fingerprint accuracy values are higher than chance levels empirically estimated from permutation analysis where subject identifiers are randomized. The plot on the left shows the IIT discovery set, while the plot on the right shows the Gi replication set. Panel D shows accuracy values (IIT discovery, left; Gi replication, right) when fingerprint accuracy is evaluated separately for each individual stimulus. These plots group accuracy values by semantic feature categories. Bars depict the mean accuracy over all stimuli within a semantic feature category, while the error bars depict ±1 standard error of the mean.

### Procedure

Eye tracking data in the IIT dataset was collected on a Tobii Pro Spectrum machine with a sampling rate of 150 Hz. Data from the Gi dataset, data was collected on an EyeLink 1000 (SR Research, Ottawa, Canada) with a sampling rate of 1 kHz. Participants sat in front of the eye tracking monitor at a distance of approximately 60 (IIT) or 65 cm (Gi) with their chin on a chinrest. Each participant completed 2 eye tracking sessions separated by an average of 7 (IIT) to 16 (Gi) days. Within each session, participants viewed all 700 stimuli, broken into 7 blocks of 100 stimuli each. Each stimulus was shown for a duration of 3 seconds. Participants were given the instructions to watch the stimuli in a natural fashion (e.g., ‘look at the images in any way [they] want’). Stimulus presentation was randomized such that within each session participants viewed the stimuli in a random order. Between the presentation of each stimulus was a fixation cross in the center of the screen. Between stimuli the IIT dataset used a jittered duration between 500 to 1000 milliseconds, whereas in the Gi dataset proceeding to the next stimulus was self-paced required a button press to move to the next stimulus. At the end of each block participants were given the opportunity to rest before starting the next block. The entire eye tracking session with all 7 blocks lasted approximately 60 minutes in the IIT dataset. Stimulus presentation was implemented within MATLAB 2019b using Psychtoolbox 3.0.16 for the IIT dataset, and MATLAB 2016b and Psychtoolbox 3.0.12 for the Gi dataset. Calibration was run using 5 (IIT) or 9 (Gi) calibration and validation points and was repeated if necessary. Within the IIT dataset, we accepted an average calibration accuracy across points and across the two eyes less than 2 degrees of visual angle. There were no differences in calibration accuracy in the IIT dataset between session 1 and session 2 (session 1: mean = 0.47 degrees, SD = 0.29 degrees; session 2: mean = 0.47 degrees, SD = 0.22 degrees; t(104) = 0.003, p = 0.99). Additionally, there were no associations between calibration data quality and gaze fingerprinting measures or other behavioral variables (e.g., autistic traits) (see Supplementary Table 1). Calibration data quality for the Gi dataset was not available and thus could not be explored further.

### Behavioral Measures

At the end of the first session in the IIT discovery dataset, participants completed the Wechsler Abbreviated Scale of Intelligence (WASI) in order to characterize full-scale IQ. At the end of the second session, participants were asked to complete two self-report questionnaires measuring autistic traits, translated into Italian - the Autism Quotient (Baron-Cohen et al., 2001) and the adult self-report form of the Social Responsivity Scale, Second Edition (SRS-2) (Constantino & Gruber, 2012). A small subset of individuals (n=9) fell above clinically-significant cut-offs of 26 on the AQ or a 2 SD cutoff from typically-developing norms of the Italian translation of the SRS-2 adult self-report (i.e. total raw score > 80.81). These individuals are annotated in figures showing autistic traits in order to visually mark individuals with or without pronounced elevation in autistic traits.

### Gaze fingerprinting analysis

Fixations were defined using velocity-based classification algorithm whereby eye movements are identified based on a threshold when the velocity of directional shifts of the eye exceed 30 degrees of visual angle per second. Data below this velocity-based threshold are considered fixations. Fixations with a duration below 100 milliseconds were excluded. These criteria for defining fixations were identical across IIT and Gi datasets. The x and y coordinates within a time-epoch defined as a fixation are then averaged to get the fixation coordinates. The duration of each fixation is the time elapsed between onset of the first gaze data sample and offset of the last data sample included in the fixation epoch. Fixation data coordinates were used to compute gaze heatmaps for each stimulus and participant. Gaze heatmaps were constructed by taking fixation points and smoothing them by a Gaussian kernel equal to a radius of 1 degree of visual angle and then normalizing the final heatmap by the maximum value (Xu et al., 2014). Heatmaps were then utilized to compute intra-subject and inter-subject gaze similarity. Gaze similarity is generally defined as the Pearson correlation between two vectorized gaze heatmaps (Kennedy et al., 2017). Intra-subject gaze similarity corresponds to the similarity value between session 1 and 2’s heatmaps for the target individual. Inter-subject gaze similarity for a particular stimulus is the between-subject similarity when the target individual’s session 1 heatmap is compared to a distractor individual’s session 2 heatmap.

Gaze fingerprinting analysis utilizes both intra-subject and inter-subject gaze similarity values. A symmetric gaze similarity matrix of size [n_subjects, n_subjects] is first computed whereby along the diagonal of this matrix are the intra-subject gaze similarity values. The off-diagonal values of this matrix represent all other pairwise inter-subject similarity values when the target individual’s session 1 heatmap is compared to the distractor individual’s session 2 heatmap. Once this matrix has been computed, we implement a ‘find-the-biggest’ (i.e. argmax) algorithm (Finn et al., 2015; Haxby et al., 2001; Kennedy et al., 2017) whereby we loop through the rows of the gaze similarity matrix and identify the subject index corresponding to the maximum gaze similarity value. A ‘hit’ is defined as a situation when the max gaze similarity value corresponds to the target individual in question. In contrast, a ‘miss’ occurs when the max gaze similarity value corresponds to some other distractor individual. Fingerprint accuracy can then be computed as the number of hits divided by the total number of participants. Null distributions for fingerprint accuracy are also computed by first permuting subject identifiers and then re-running the fingerprint analysis pipeline 1000 times. From these null distributions, p-values can be computed by counting the number of times fingerprint accuracy was as large or larger than the real fingerprint accuracy and then dividing that number by 1001.

Gaze fingerprinting analysis, as defined above, was applied at-scale to every stimulus. Additionally, we also applied gaze fingerprinting analysis to averaged similarity values across all or subsets of the stimuli set grouped by semantic features. To mimic the gaze fingerprinting analysis procedure of Kennedy et al., (Kennedy et al., 2017), we first computed median gaze similarity by computing the median over all 700 stimuli. The median was chosen to better reflect central tendency of the distribution than the mean, particularly in situations where the distribution is skewed. When distributions are not heavily skewed, the mean and median should be relatively similar. With a 3-dimensional gaze similarity matrix of size [n_subjects, n_subjects, n_stimuli], we compute the median over the 3^rd^ dimension of the 700 stimuli to get a median gaze similarity matrix of size [n_subjects, n_subjects]. The fingerprint analysis algorithm described above is then used to define ‘hits’ or ‘misses’ and compute accuracy. This same procedure of computing a median gaze similarity matrix was also redone on subsets of stimuli that correspond to each of the 12 semantic features labelled in the OSIE dataset (Xu et al., 2014). To get 95% confidence intervals (CI) around accuracy when averaging all stimuli, we utilized bootstrapping with 10,000 resamples. This allowed us to compare accuracy rates within semantic feature categories to the lower bound bootstrap CI on accuracy computed when averaging across all stimuli.

The ‘hit’ or ‘miss’ nature of the gaze fingerprinting output is a binary output that may hide information regarding the continuous nature of ‘fingerprintability’ or degree of ‘gaze uniqueness’. That is, the fingerprint output may be a ‘miss’ because the intra-subject similarity is not the maximum gaze similarity among all distractor individuals. However, in the instance where the intra-subject similarity is still ranked as very large (e.g., 2^nd^ highest gaze similarity), this would be a situation whereby the individual’s gaze is unique to some high degree, though not maximally unique when compared to all distractor individuals. Thus, to derive a continuous metric of degree of ‘gaze uniqueness’, we developed a score we call the gaze uniqueness index (GUI) that conveys in a continuous manner the degree of gaze uniqueness that would otherwise be lost in a binary fingerprint output of hits or misses. GUI is computed by dividing intra-subject similarity by average inter-subject similarity. The denominator in this equation of average inter-subject similarity is important for interpretational reasons, as it acts to scale the intra-subject similarity to what can be thought of as a ‘norm’ or ‘baseline’ level of similarity for a particular stimulus, represented by the average of all pairwise inter-subject similarities for the target individual. Given that GUI values are ratios of intra-subject/average inter-subject similarity, the null value is 1, indicating equal intra-subject and average inter-subject gaze similarity. Values significantly increased greater than 1 signify a global gaze pattern across all stimuli that is relatively unique to the individual. GUI values are computed at-scale per all 700 stimuli and are also summarized as one number per participant by taking the median GUI value across all 700 stimuli. Hypothesis tests on GUI values per each subject are implemented with a one-sample t-test against a null value of 1. From these t-tests, p-values are computed and then we use false discovery rate (FDR) multiple comparison correction to identify individuals that pass FDR q<0.05.

### Gaze fingerprint barcoding

Given the stimulus-rich nature of the OSIE dataset (e.g., 700 stimuli), gaze fingerprinting at-scale for each subject and stimulus allows for an [nsubs, nstimuli] binary output matrix of hits and misses. The rows of this matrix can be thought of as individualized barcodes that tell us which stimuli are fingerprintable for that individual. Given that there is no requirement for each individual’s barcode to be similar or different, it is an open question as to whether such gaze fingerprinting barcodes are unique to individuals or are clustered into subsets of individuals with similar kinds of barcodes. To answer this question, we compute a Jaccard distance matrix to represent the dissimilarity between gaze fingerprinting barcodes and then utilize hierarchical clustering (ward.D2 agglomeration method) on this distance matrix to identify how many clusters are represented in the dataset. This analysis allows us to answer what the optimal number of clusters (k) is using the silhouette coefficient over a range of k=2 up to k = n-1 as the maximum number of clusters. In a situation where k = n-1, such a solution would indicate that each individual’s gaze fingerprint barcode is unique and that clusters are individual subjects.

Gaze fingerprint barcodes can also be described by enrichment with semantic feature categories. To achieve these goals, we computed enrichment odds ratios and p-values in a manner identical to an analogous statistical procedure in bioinformatics known as gene set enrichment analysis. In this type of analysis, we examine the extent of overlap between two lists (e.g., fingerprintable stimuli and semantic feature stimuli) whereby effect size is indicated by an odds ratio. This enrichment odds ratio can be interpreted with respect to the null value of 1, whereby values increasingly higher than 1 indicate an over-representation of a specific semantic category within an individual’s fingerprintable stimuli set. To compute odds ratios, we first compute a 2×2 contingency table, with cells marked A and B in row 1 and cells C and D in row 2. Counts in cell A reflect the number of fingerprintable stimuli given a semantic feature. Counts in cell B are the number of fingerprintable stimuli minus the number of fingerprintable stimuli given a semantic feature. Counts in cell C are the total number of stimuli within a semantic feature category minus the number of fingerprintable stimuli given a semantic feature. Finally, counts in cell D correspond to the total number of stimuli in the OSIE dataset minus the summed quantity in cells A, B, and C. Such a contingency table can be inserted into the *fisher.test* function in R and get as output an enrichment odds ratio. These enrichment tests were computed per each subject using the subset of fingerprintable for each subject and testing the overlap of that stimulus set with the stimuli within each of the 12 semantic categories. A resulting matrix of [nsubs, nfeatures] can be computed as output of this analysis and then assessed for subtypes via hierarchical clustering analysis, where optimal k is identified by majority vote amongst many clustering metrics computed in the NbClust package in R (Charrad et al., 2014). This analysis allows for discovery of subtypes characterized by different patterns of semantic feature enrichment.

### Association analysis with autistic traits

Robust regression linear modeling was utilized as the modeling strategy to test for association between gaze fingerprinting metrics and autistic traits. Autistic traits were considered the dependent variable in the model. Autistic traits were initially measured via two different questionnaires, the AQ and SRS-2. These measures are highly correlated (r = 0.77, p < 2.2e-16), indicating that shared variance in both may best account for autistic traits as a latent variable. To isolate the shared component of variation in AQ and SRS-2, we utilized principal components analysis (PCA) and found that the first principal component accounts for 88% of the shared variance. This autistic traits PC1 was used as the dependent variable measuring autistic traits. Two gaze fingerprinting metrics were utilized as independent variables in the model – 1) fingerprintable stimuli count and 2) median gaze uniqueness index computed across all 700 stimuli. Rather than plugging both variables into the model as separate independent variables, we first checked to see if the two fingerprinting metrics were correlated and found that they were (r = 0.81, p < 2.2e-16). Thus, we instead used PCA to decompose fingerprinting metrics into two orthogonal variables, of which PC1 captured around 91% of the variance and loaded equally onto both metrics, while PC2 accounted for the remaining 9% of the variance. With PC1 and PC2 gaze fingerprinting metrics, these variables were utilized as independent variables in the linear model to test for associations with autistic traits. Finally, the other independent variable used as a covariate of no interest was sex. This was done because prior research has shown robust sex differences in autistic traits on both AQ and SRS-2 (Baron-Cohen et al., 2001; Constantino & Todd, 2003; Frazier et al., 2014). Thus, our modeling strategy uses sex as a covariate before assessing the association of fingerprinting metrics with autistic traits. The linear model was a robust regression linear model and was utilized to guard against the influence of bivariate outliers and was implemented with the *lmrob* function in the *robustbase* library in R. The *lmrob* function was also utilized to draw best fit lines in scatterplots to visualize the results.

We also compared 3 different models predicting autistic traits PC1. All of the models first accounted for sex as a covariate. Model 1 utilizes gaze fingerprinting PC1. Model 2 utilizes the median intra-subject gaze similarity computed across all 700 stimuli. Finally, model 3 utilizes the median average inter-subject gaze similarity computed across all 700 stimuli. Models 2 and 3 can be thought of as ‘null’ models with respect to the model of importance for gaze fingerprinting (e.g., model 1). Since gaze fingerprinting metrics (i.e. number of fingerprintable stimuli, median GUI) are computed based on intra-subject and inter-subject similarity values, comparing model 1 to models 2 and 3 allow us to assess the importance of gaze fingerprinting metrics relative to models that predict autistic traits solely on the basis of components used within gaze fingerprinting – that is, intra-subject similarity (model 2) or inter-subject similarity (model 3). Percentage of variance explained (R^2^) was computed for each model and compared to each other, with the better model being the model that explains more variance (e.g., higher R^2^) in autistic traits PC1.

### Pre-registered short-term and long-term delay follow-up experiments

The gaze fingerprint barcodes reported from the Discovery dataset are derived from eye tracking sessions 1 and 2, separated by about 7 days. With only one barcode per individual, it could be claimed that the barcodes derived from the Discovery dataset are essentially a random sequence of ‘hits’ and ‘misses’. If such a situation were true, it would lead to a hypothesis that measuring barcodes on the same individual in further follow-up repeat eye tracking sessions would lead to barcodes that are not similar to the original barcodes derived from the original sessions 1 and 2. To test this null hypothesis that gaze fingerprint barcodes are random and not significantly similar within-individual, we embarked on 2 follow-up pre-registered experiments where we measured gaze fingerprint barcodes within-individual after either short-term (19-90 days) or long-term (∼ 1.87-5.16 years) delays and then evaluate whether those later follow-up barcodes are significantly similar to the original barcodes derived from individuals in the original sessions 1 and 2. The long-term follow-up experiment included only one additional eye tracking session after the long delay period on the order of years. This allowed for one comparison of barcode similarity between sessions 1-2 (7 day separation) and sessions 1-3 (1.87-5.16 year separation). In contrast, the short-term follow-up experiment was designed to have 5 repeat eye tracking sessions, which allowed for 3 repeated measurements of barcodes and for similarity between session 1-2 barcodes (7 day separation) to be compared with barcodes for sessions 1-3, 1-4 and 1-5 (19-90 days separation). While the long-term delay experiment allowed for only one measurement of a barcode and thus, does not allow for multiple repeat barcodes to be derived after such long-term delays, the 5 repeat eye tracking sessions included in the short-term follow-up experiment allowed for examination of barcode similarity within-individual as a function of time on the order of months. This unique longitudinal characteristic of the short-term follow-up experiment allows for tests of how barcode similarity changes (or not) over time within an individual. For the pre-registration on OSF please see: https://osf.io/atj7b. Below we will explain the methods and analysis behind these follow-up experiments.

For the experiment testing barcode similarity over long-term delays, we aimed to test n=10 participants from the original Discovery dataset in a single eye tracking ‘session 3’ that was done after an approximate 5-year delay from their original eye tracking sessions 1 and 2. After applying identical data quality filtering as in the original Discovery dataset experiment (i.e. including only individuals with missing gaze data in less than 15% of stimuli), these n=10 individuals were retained for further analysis. These individuals repeated the same experimental paradigm and procedure as sessions 1-2 (e.g., viewing 700 static pictures of complex natural scenes from the OSIE stimulus set). Eye tracking session 3 from these individuals was separated by an average of 3.58 years (SD = 0.93 years, range = 1.87-5.16 years) from the original eye tracking sessions 1 and 2 and allowed us to compute a gaze fingerprint barcode for these individuals using data from sessions 1 and 3. The analysis pipeline to compute gaze fingerprints and gaze fingerprint barcodes was identical to the Discovery experiment and all individuals from the original Discovery dataset were utilized as distractor individuals within the gaze fingerprint classifier. With two gaze fingerprint barcodes derived from each individual in the long-term follow-up experiment – that is, one barcode derived from sessions 1 and 2 separated by approximately 7 days and another barcode derived from sessions 1 and 3 separated by 1.87-5.16 years – we can test the null hypothesis that over long periods of time (e.g., years) gaze fingerprint barcodes are random and not a repeatable individuating code that signifies what is unique about what an individual looks at within static pictures complex natural scenes. To quantify the similarity between session 1-2 and session 1-3 barcodes, we computed Jaccard distance between these two barcodes, identical to the similarity metric used in the original Discovery dataset clustering analyses. We then compared the real barcode similarity to a distribution of Jaccard distance values simulated under the null hypothesis by permuting the session 1-3 barcode and the re-computing Jaccard distance with session 1-2’s barcode. This permutation analysis was run 10,000 times to build the null distribution of Jaccard distance between barcodes. A p-value can then be computed on a within-individual basis as the number of times barcode similarity was as low or lower than the real barcode similarity, divided by 10,001. Such hypothesis tests were done on all n=10 individuals in the long-term follow-up experiment. We thresholded each individual’s test at an alpha level of 0.05 to get a count of the number of individuals within the long-term follow-up experiment with significantly similar barcodes after the long-term delay. This count was then used in a binomial test to test whether the number of identified significant individuals deviates from the number of significant individuals that would be expected under the null hypothesis.

The second pre-registered follow-up experiment was conducted to examine barcode similarity over short-term delays of 19-90 days. This was done as a contrast to what occurs in the long-term delay experiment, but also to examine within-individual longitudinal change in barcode similarity over time. In this short-term follow-up experiment we aimed to test n=10 individuals over 5 repeat eye tracking sessions that were identical to those in the original Discovery and long-term delay experiments. Although the aim was to collect n=10 individuals, one participant was dropped due to lack of data on return sessions 3-5. Furthermore, after applying data quality filtering identical to the original Discovery dataset experiment (i.e. including only individuals with missing gaze data in less than 15% of stimuli), the final sample sizes were n=7 for the delay between sessions 1-3, n=6 for the delay between sessions 1-4, and n=7 for the delay between sessions 1-5. The delay between sessions 1 and 2 were similar to the original Discovery dataset (e.g., mean = 7.80 days, SD = 1.47 days, range = 7-11 days). Session 3 was separated from session 1 by a delay of 25.43 days on-average (SD = 5.89 days, range = 19-36 days), while session 4 was separated by 36 days on-average (SD = 1.47 days, range = 35-38 days), and session 5 was separated by 72 days on-average (SD = 12.31 days, range = 56- 94 days). In terms of analysis, we conducted identical permutation tests within-individual and binomial test of counts of significantly individuals, as described above for the long-term experiment. We additionally ran further longitudinal modeling, given that each subject had multiple repeat measurements of barcodes of time. The variables used for longitudinal modeling were time delay from session 1 measured in days and real barcode similarity (i.e. Jaccard distance between barcodes for session 1-2 and session 1-3) versus mean permuted barcode similarity. This distinction between real versus mean permuted barcode similarity was a factor called ‘condition’ in the model. Longitudinal modeling was implemented with linear mixed effect models (e.g., the *lmer* function within the *lmerTest* library in R), whereby the dependent variable was barcode similarity. The fixed effect independent variables in the model were time (i.e. number of days delay since session 1) and condition (i.e. real versus permuted) and the time*condition interaction, while the random effect of time was modeled within-individual with random intercepts and slopes. The main effect of time allows for examination of whether barcode similarity changes as a function of the delay between session 1. The main effect of condition allows for a test of whether barcode similarity is significantly different under real versus permuted conditions. The interaction between time and condition allows for a test of whether barcode trajectories over time significantly differ from each other.

### Data and code availability

All code and data for reproducing analysis can be found within our GitHub repository: https://github.com/IIT-LAND/gaze_fingerprint_osie.

## Results

In our first analysis we tested whether gaze fingerprinting is possible, and if so, how good is it at accurately detecting individuals as a function of semantic features within stimuli. As a first approach to answering these questions, we averaged gaze similarity across stimuli. Here we found fingerprint accuracy of 52.40% in the IIT discovery dataset and 62.98% in the Gi replication dataset (Figure 1B). Both accuracy rates are well above the theoretical (e.g., ∼1- 2%) and empirically-derived (via permuting subject identifier) chance accuracy rate (p < 9.99e- 4). In sex-stratified analysis on the IIT discovery dataset, sample sizes are comparable to the Gi dataset, and the accuracy rates fall within a similar range of approximately 48-65% (e.g., 48% in females, 65 % in males) (see Supplementary Figure 1).

Split into different subsets of stimuli reflecting high-level semantic features, accuracy in all features are well above chance levels in both discovery and replication sets (IIT: 29-54%, all p < 9.99e-4; Gi: 39-63%, all p < 9.99e-4) (Figure 1B). Notably, accuracy levels for each semantic feature are within the 95% bootstrap confidence intervals computed when averaging across all stimuli in both IIT and Gi datasets. The exceptions to this are smell, sound, emotion, and animal features. These features are the most infrequently occurring stimuli (e.g., comprising <22% of all stimuli) (e.g., Figure 1A). Thus, relatively lower accuracy on these features is likely due to number of stimuli averaged within the semantic feature set. In summary, the relatively high accuracy rates when averaging across all stimuli can also be achieved by averaging across stimuli within semantic features, provided that the semantic feature category contains an adequate number of stimuli (e.g., >154 stimuli).

Given that accuracy tends to drop with number of stimuli present within a semantic feature category, it is likely that identification accuracy applied at-scale to each stimulus would result in relatively low accuracy. However, would identification accuracy computed per each stimulus be higher than chance identification rates? Indeed, we find that identification accuracy at the level of individual stimuli is heavily reduced compared to accuracy on averaged gaze similarity (e.g., IIT: ∼2-11%; Gi: ∼4-21%) (Figure 1C). Average accuracy of stimuli grouped by semantic features also shows attenuated accuracy at the individual stimulus level (IIT: 3.9- 5.1%; Gi: 5.2-6.6%) (Figure 1D). However, we can evaluate how many of the 700 stimuli show accuracy rates significantly above chance at FDR q<0.05. Here we find in the IIT discovery set that around 94.2% (660/700) of stimuli allow for above chance accuracy. This percentage was much lower for the Gi replication set (62.2%; 436/700) (Figure 1C). These results generally suggest that gaze fingerprinting is possible and that averaging across a range of different stimuli can enhance detection rate. While most individual stimuli show statistically higher than chance levels of accuracy, accuracy for individual stimuli is far lower than averaging gaze similarity across stimuli. This suggests that gaze fingerprinting based on any one stimulus in isolation may not achieve high levels of accuracy across a population.

While the accuracy of around 52-63% for averaged gaze similarity across stimuli is already impressive and much higher than previous work on twins or with a small number of movie stimuli (e.g., ∼29-40% accuracy (Keles et al., 2022; Kennedy et al., 2017; Rigas & Komogortsev, 2014)), it is possible that averaging across stimuli may hide important individuating information within the patterning of fingerprintability across stimuli. Given the unique stimulus-rich characteristic of our dataset, we next utilized a ‘gaze fingerprint barcoding’ approach to test whether the patterning of fingerprintability across individual stimuli would allow for more individuation and fingerprint detectability in all participants. The gaze fingerprint barcoding approach uses the binary vector of hits or misses across all 700 stimuli as a potentially unique individuating gaze fingerprint pattern for an individual. As shown in Figure 2C, gaze fingerprint barcodes are relatively sparse due to each individual showing a small, but potentially unique pattern of fingerprintable stimuli across all 700 OSIE stimuli. To begin teasing apart whether this approach could yield further insights beyond those gleaned from fingerprinting based on averaged gaze similarity, we first ask whether the sparse information within barcodes, that is - the number of fingerprintable stimuli detected per individual, is larger than what we would expect at chance when subject identifier is randomly permuted. Here we find that number of fingerprintable stimuli varies widely between individuals, with some individuals showing very minimal levels of fingerprintable stimuli to others who can be identified based on 12-14% of all stimuli (Figure 2A). We replicably find that nearly all individuals (IIT: 96/105 or 91.4%; Gi: 44/46 or 95.6%) had fingerprintable stimuli counts that were significantly higher than chance when subject identifier is randomly permuted (Figure 2A). Thus, the information present within sparse gaze fingerprint barcodes is non-random and much higher than would be expected at chance when subject identifier is randomly permuted.

**Figure 2:**
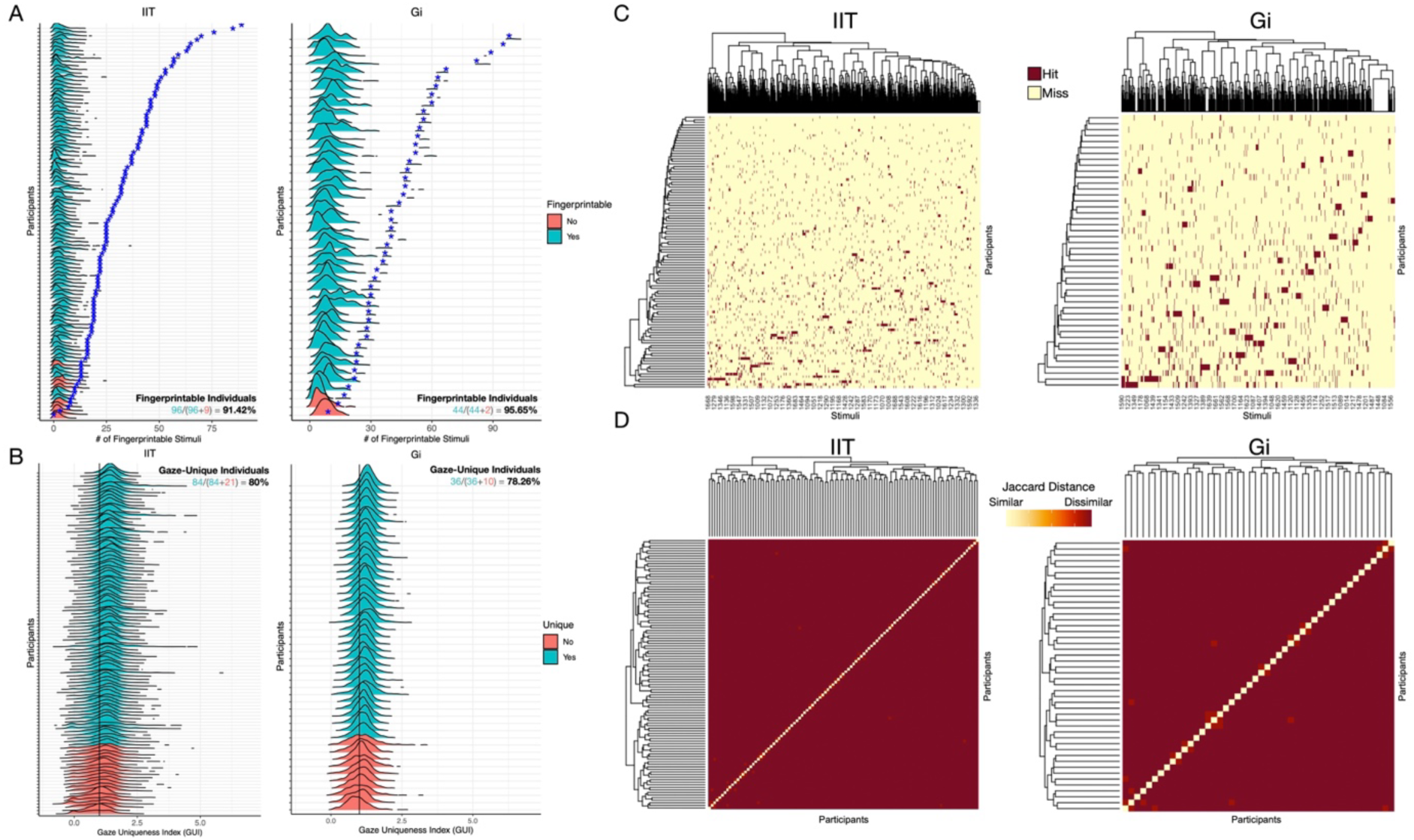
Gaze fingerprint barcoding and degree of gaze uniqueness. Panel A shows counts of fingerprintable stimuli (x-axis) per each individual (y-axis) (IIT discovery, left; Gi replication, right). The actual count for fingerprintable stimuli per subject is denoted by the blue star. The density plots per each individual represent the null distribution of fingerprintable stimuli count over 1000 permutations where subject identifier is randomly permuted. The colors denote a distinction of whether the individual is fingerprintable (turquoise) or not (pink), with turquoise individuals having an actual fingerprintable stimuli count (blue star) that is statistically significantly elevated relative to their null distribution (density plots) and at a false discovery rate (FDR) of q<0.05. As annotated on the bottom right of the plots, ∼91% of IIT discovery and ∼95% of Gi replication subjects are fingerprintable in having statistically significant fingerprintable stimuli counts. Panel B depicts a continuous metric indicative of the degree of gaze uniqueness per individual (IIT discovery, left; Gi replication, right). The gaze uniqueness index (GUI) shown on the x-axis is computed as the ratio between intra-subject divided by average inter-subject gaze similarity. Increasing GUI values indicates increasing gaze uniqueness, whereas values near 1 indicate equal levels of intra-subject and average inter-subject gaze similarity. The density plots per each subject shown on the y-axis depict GUI values across all 700 stimuli, and statistical tests were computed per each individual to test whether GUI was significantly elevated relative to the null value of 1 (vertical black line). After FDR correction for multiple comparisons, subjects can be colored according to individuals that have GUI values significantly different from 1 (turquoise) versus those that do not significantly differ from the null value of 1 (pink). Panel C shows a visual representation of ‘gaze fingerprint barcode’ matrices (IIT discovery, left; Gi replication, right). These matrices have subjects along the rows and stimuli along the columns and depict binary values that indicate ‘hit’ (dark red) or ‘miss’ (light yellow) output of the gaze fingerprint classifier. A ‘hit’ indicates that an individual can be fingerprinted for that particular stimulus. A ‘miss’ indicates that the individual cannot be fingerprinted for that particular stimulus. Panel D represents between-subject Jaccard dissimilarity matrices computed on binary gaze fingerprint barcode matrices. The diagonal represents an individual’s barcode similarity with themselves and is maximially similar (e.g., light yellow). The off-diagonal values depict how similar/dissimilar are gaze fingerprint barcodes between individuals. Rows and columns of this matrix are re-arranged according to hierarchical clustering dendrograms, such that subtypes would appear as blocks along the diagonal. The absence of such subtype blocks along with near maximal dissimilarity for all between-subject comparisons illustrates visually what clustering algorithms identify as the optimal clustering solution of k = n-1. This solution indicates that each individual is its own cluster. This result illustrates the uniqueness represented in the sparse information conveyed by gaze fingerprint barcodes.

As an alternative to a binary vector representing the gaze fingerprint barcode, we introduce the concept of degree of ‘gaze uniqueness’, via a metric we call the ‘gaze uniqueness index’ (GUI). GUI can be computed on a per-stimulus basis and is defined as the ratio of intra-subject gaze similarity divided by average inter-subject gaze similarity. Increasing GUI values indicate increasing degree of gaze uniqueness, whereas the null value for GUI of 1 indicates equal intra-subject compared to average inter-subject gaze similarity. Hypothesis tests were done on the set of all 700 stimuli to test whether an individual has a GUI significantly above 1, indicating a relatively unique gaze profile on-average across all stimuli. As shown in Figure 2B, about 80% of IIT (84/105) and 78.2% (36/46) of Gi individuals show a GUI significantly greater than the null value of 1. Thus, GUI can also capture individuating information for a sizable majority of individuals but may not be as sensitive as the sparser signal present in gaze fingerprint barcodes.

Gaze fingerprint barcodes (Figure 2C) can also be utilized within a clustering framework to identify whether barcodes are unique to an individual or whether they cluster together in subsets of similar types of individuals. Using Jaccard distance to quantify how similar or dissimilar the binary gaze fingerprint barcodes are between-individuals, we can then apply clustering analysis to this between-subject distance matrix and identify how many clusters exist. If barcodes are unique individuating codes, clustering should identify that the maximum number of clusters detectable (e.g., k = n-1) will be the optimal number of clusters. Indeed in both discovery and replication datasets, we find that the optimal number of clusters is k = n-1 (Figure 2D). This result confirms that gaze fingerprint barcodes hold unique individuating information and that such insights can only be gleaned by gaze fingerprint profiling and barcoding of individuals at-scale across a large set of stimuli.

An alternative explanation for this clustering result on gaze fingerprint barcodes could be that the barcodes are random. To test this would require further follow-up experiments on the same individuals to measure barcodes repeatedly and then assess their similarity. Therefore, to test this hypothesis, we conducted two pre-registered follow-up experiments. In the first follow-up experiment, we called back n=10 of the original individuals tested in the Discovery experiment for a 3^rd^ eye tracking session where they repeated the same paradigm as in sessions 1 and 2. However, this 3^rd^ eye tracking session was conducted after years of delay compared to session 1 (e.g., mean delay 3.58 years, SD = 0.93 years, range = 1.87-5.16 years). Long-term gaze fingerprint barcodes were computed by comparing gaze heatmaps from session 1 to session 3 and then running the same gaze fingerprint classifier as utilized in the original Discovery experiment. For these n=10 individuals, we could then compare gaze fingerprint barcodes derived from the original Discovery experiment (e.g., delay of 7 days between sessions 1 and 2) to the barcodes derived from the long-term follow-up where the delay between session 1 and 3 is on the order of years. Permutation tests on Jaccard distance between gaze fingerprint barcodes were computed for each of the n=10 individuals to identify the number of individuals where gaze fingerprint barcode similarity is significantly different from chance similarity when barcodes are randomizes. Of the n=10 individuals tested, 30% had significantly similar barcodes relatively to chance and a binomial test identifies this as statistically significant number of successful hypothesis tests (number of successes = 3, number of trials = 10, p = 0.0115). A follow-up correlation test between barcode similarity and delay between sessions 1 and 3 revealed no correlation between time delay and barcode similarity (r = −0.15, p = 0.67). Thus, this result suggests that gaze fingerprint barcodes are stable over long periods on the order of years for a significant number of individuals, and that barcode similarity is not associated with the length of time between compared barcodes.

We next conducted another pre-registered follow-up experiment whereby individuals were gaze tracked over 5 repeat sessions separated by 19-90 days, as a contrast to tests of barcode similarity on the time scale of years (e.g., the long-term follow-up experiment). Barcodes computed from sessions 1 and 2 were compared to barcodes from sessions 1-3, 1-4, and 1-5, with a similar permutation and binomial test approach. Compared to barcodes from sessions 1-2, 71% of individuals had significantly similar barcodes in sessions 1-3 (number of successes = 5, number of trials = 7, p-value = 6.027e-06), while 33% were significantly similar in sessions-14 (number of successes = 2, number of trials = 6, p-value = 0.03277), and 28% were significantly similar in sessions 1-5 (number of successes = 2, number of trials = 7, p-value = 0.04438). This data can also be evaluated with longitudinal modeling, given the repeat assessments of barcode similarity within-individual over time. Figure 3 shows that there is no significant effect over time (F = 1.34, p = 0.29). Comparing real barcode similarity to mean permuted barcode similarity, there is a significant effect whereby real barcode similarity is significantly more similar that barcode similarity under null hypothesis conditions (e.g., randomized barcodes) (F = 13.58, p = 0.001). Finally, there is no interaction between time and condition (F = 2.23, p = 0.13), indicating that the trajectories of barcode similarity under real versus permuted conditions are not significantly different from each other.

**Figure 3:**
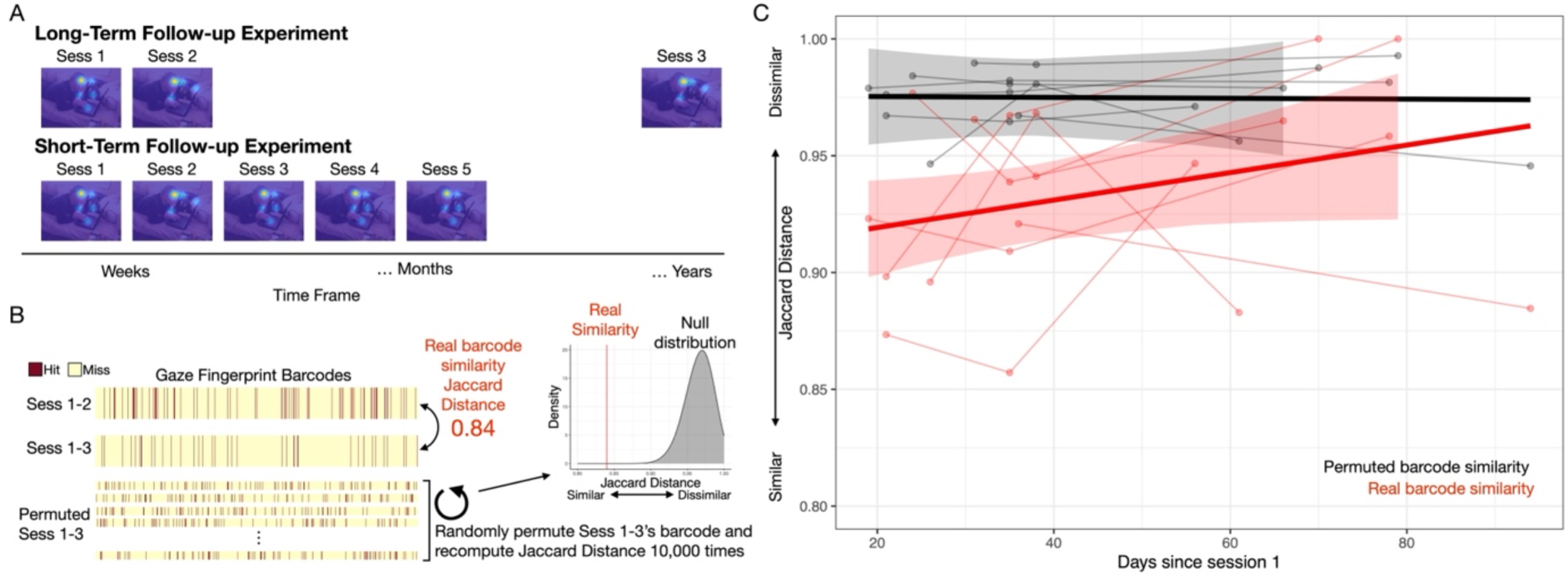
Stability of gaze fingerprint barcodes over time. Panel A shows a schematic of the design of the pre-registered long-term and short-term follow-up experiments whereby we tested individuals after a delay from session 1 on the order of years (e.g., long-term) or short-term (e.g., weeks to months). The short-term follow-up experiment had 5 repeat eye tracking sessions, which additionally allowed for tracking of gaze fingerprint barcodes within-individual over the course of 2-3 months. Panel B shows a schematic of the data analysis approach for assessing similarity of gaze fingerprint barcodes. Jaccard distance was utilized to quantify degree of barcode similarity, with smaller Jaccard distance being indicative of more barcode similarity. Barcodes were then randomly permuted 10,000 times to build a null distribution of Jaccard distance under the hypothesis that the barcodes are random. Statistical inference can be implemented on a within-individual level by computing a p-value that describes the probability of getting a barcode similarity as small or smaller than the real barcode similarity in unpermuted data. In the example show, the real barcode similarity is Jaccard distance = 0.84. Under the simulated null distribution, such barcode similarity is never observed and hence the p-value is 1/10001 = 9.99e-5. Panel C shows barcode trajectories in the short-term follow-up experiment for real (red) and mean permuted (black) conditions. Each individual is a line and the group trajectory is plotted in the thicker solid line with 95% confidence bands. Time since session 1 in days is plotted on the x-axis, while barcode similarity (Jaccard distance) is plotted on the y-axis, with smaller values indicating more barcode similarity.

Overall, these results under short-term follow-up time frames on the order of months match the inferences from the long-term follow-up – that is, gaze fingerprint barcodes are significantly similar for a significant number of individuals. Furthermore, the lack of a correlation between time delay and barcode similarity in the long-term follow-up along with the lack of a significant effect of time in the longitudinal modeling of the short-term follow-up indicates that barcodes are highly stable individualized traits that are not susceptible to large degrees of change over time.

Since the gaze fingerprint barcode is a stable non-random individuating sequence of information, we next asked whether there is any systematic pattern of semantic feature enrichment within an individual’s fingerprintable stimuli set. To answer this question, we went back to the original Discovery dataset and computed semantic feature enrichment odds ratios that confer an effect size relevant to how many fingerprintable stimuli fall within a particular semantic category. These semantic feature enrichment odds ratios can be thought of as ‘semantic feature enrichment barcodes’ that semantically describe each individual’s fingerprintable stimuli set. Thus, we applied a similar clustering approach to assess how unique are these types of barcodes amongst individuals. Here we find replicable evidence of two clusters of participants in both discovery and replication datasets (Supplementary Figure 2A-B). One of these subtypes comprises a small minority of the sample (IIT: 13.5%; Gi: 6.8%) and can be described as individuals whose fingerprintable stimuli are largely enriched for primarily social features of stimuli (e.g., social, humans, faces) and with an under-representation of non-social features (Supplementary Figure 2C-D). However, the majority of individuals in IIT and Gi samples fell within a cluster where there was no unique signature with respect to enrichment in semantic features. These results suggest that stratification of the population based on the over-representations of specific semantic features within fingerprint barcodes may be limited to a small subset of individuals.

In a final set of analyses, we expanded scope on the translational relevance of gaze fingerprinting to autistic traits. Autistic traits represent a continuous and highly heritable phenotype linked back to polygenic or even omnigenic common genetic architecture (Constantino & Todd, 2003; Robinson et al., 2016). Degree of gaze similarity is also strongly linked to genetic similarity (Constantino et al., 2017; Kennedy et al., 2017), and atypical gaze patterns are key markers linked to autism (Constantino et al., 2017; Jones & Klin, 2013; Wang et al., 2015). Thus, we reasoned that variation in autistic traits as a genetically sensitive phenotype may be linked to individualized patterns of gaze captured by the gaze fingerprinting approach. Here we used two self-report questionnaires that are widely used to measure autistic traits - the SRS-2 and AQ. These measures are highly correlated (r = 0.77, p < 2.2e-16) (Figure 4A) and thus, we utilized PCA to capture the first principal component accounting for about 88% of the variance shared between AQ and SRS-2. This autistic traits PC1 was our dependent variable in all further models. Gaze fingerprinting predictors in our model were a combination gaze fingerprinting metrics such as the number of fingerprintable stimuli and median GUI. However, because these variables are highly correlated (r = 0.81, p < 2.2e-16) (Figure 4B), we orthogonalize them with PCA and use PC1 and PC2 in the model as independent variables. Finally, sex was modeled as a covariate since prior research has shown strong sex differences in autistic traits in the typically-developing population (Baron-Cohen et al., 2001; Constantino & Todd, 2003; Frazier et al., 2014). Here we find that gaze fingerprinting PC1, reflecting the shared variance between number of fingerprintable stimuli and median gaze uniqueness index, is the primary significant predictor of autistic traits (β = −0.35, SE = 0.06, t = −5.16, p = 1.19e- 6) (see Supplementary Table 2). The relationship between GUI and autistic traits is a negative relationship, whereby increasing autistic traits are observed in individuals with lower gaze fingerprintability (Figure 4C). Importantly, the individuals that heavily deviate from this relationship tend to be individuals with remarkably high autistic traits, beyond cutoffs for typically-developing adult norms on SRS-2 or AQ (turquoise dots in Figure 4C). Similar results appear when AQ and SRS are analyzed separately as dependent variables (see Supplementary Table 2).

**Figure 4:**
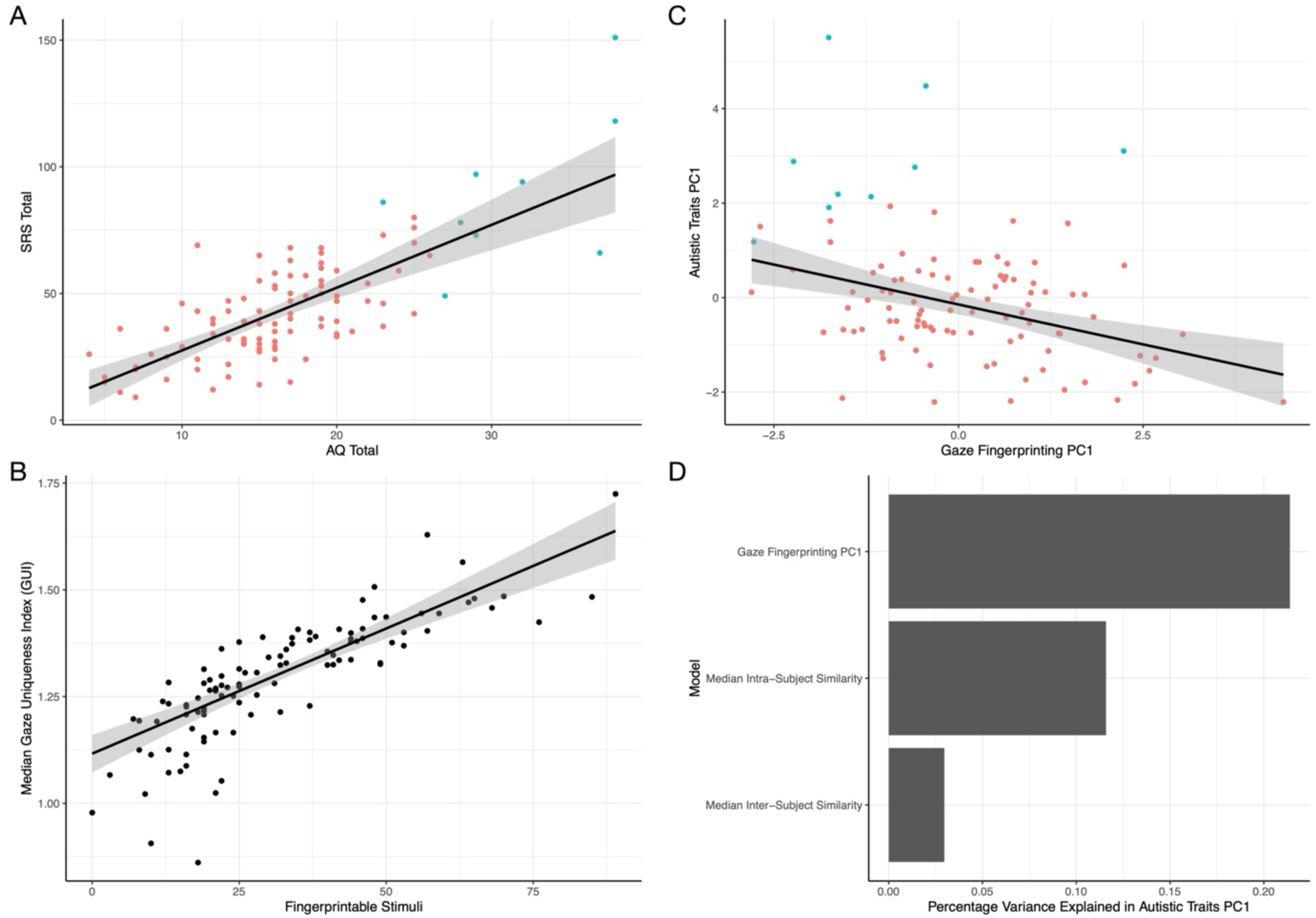
Gaze fingerprinting association with autistic traits. This figure showcases how gaze fingerprinting metrics are associated with variation in autistic traits. Panel A shows the two autistic trait measures utilized in this study – the AQ (x-axis) and SRS-2 (y-axis). SRS-2 and AQ are highly correlated and thus PCA is used to isolate the first PC representing 88% of the variance shared between AQ and SRS-2. This measure, called ‘Autistic Traits PC1’, is used as the dependent variable in our model and is plotted on the y-axis in panel C. The coloring of the dots in panel A represents two classes of individuals – individuals within the normative range expected for AQ and SRS-2 in typically-developing adults (n=96, pink) and individuals that are highly deviant from typically-developing adults (n=9, turquoise) (e.g., >26 on the AQ (Woodbury-Smith et al., 2005) or >2SD (80.81) from typically developing norms on the SRS-2). Panel B shows the 2 gaze fingerprinting measures utilized in the model – that is, the number of fingerprintable stimuli per subject (x-axis) and the median gaze uniqueness index (GUI) computed across all 700 stimuli (y-axis). The scatterplot in panel B shows that these two measures are highly correlated. Thus, PCA was used to orthogonalize the data as two principal components to use as independent variables in a linear model predicting autistic traits. Gaze fingerprinting PC1 accounts for 90% of the variance in the gaze fingerprinting metrics. Panel C shows a scatterplot depicting the association between autistic traits PC1 and gaze fingerprinting PC1. Individuals that are highly deviant from typically-developing norms are plotted in turquoise. The best fit line and confidence bands are computed from a linear robust regression model on all individuals in pink and turquoise. Panel D plots percentage of variance explained in autistic traits PC1 (x-axis) for 3 different models labeled on the y-axis. All 3 models utilize sex as a covariate and primarily differ in terms of the primary independent variable of interest, which is labeled on the y-axis. The model explaining more variance is interpreted as the better model. These model comparisons directly show that gaze fingerprinting PC1 is better at explaining variance in autistic traits compared to utilizing only information about intra-subject or inter-subject gaze similarity alone.

Next, we compared models that all incorporate sex as a covariate but differentially utilize either the gaze fingerprinting PC1 as the independent variable or use only median intra-subject or inter-subject gaze similarity. These model comparisons were done to understand whether the variance predicted by gaze fingerprinting PC1 is more than what could be predicted by only using intra- or inter-subject gaze similarity alone. Here we find that the model using gaze fingerprinting PC1 accounts for 21.39% of the variance in autistic traits, whereas median intra-subject gaze similarity accounts 11.59% of the variance and median inter-subject similarity predicts 2.96% of the variance (Figure 4D). Thus, in this direct model comparisons, it is clear that gaze fingerprinting PC1 provides significantly enhanced utility in predicting autistic traits over and above models that only take into account intra- or inter-subject similarity separately.

## Discussion

In this work we discovered that identification of individuals is possible simply by using gaze patterns sampled from 3-second viewing epochs on a large and diverse range of static visual stimuli depicting complex natural scenes. By sampling gaze patterns across 700 stimuli, we replicably show that we can gaze fingerprint individuals at levels much higher than chance (52-63% vs 1-2% chance levels). While these accuracy rates are much higher than those of previous studies (e.g., 29-40%) (Keles et al., 2022; Kennedy et al., 2017; Rigas & Komogortsev, 2014), there may be some caveats to comparing accuracy across studies. First, none of the prior studies had a stimulus-rich sampling of features similar to the current study. Therefore, it is possible that sampling across a very large array of stimuli will affect gaze fingerprinting accuracy. Indeed, we see that accuracy can drop when subsets of stimuli are not so large in size (Figure 1D). Thus, the more information sampled about how an individual looks at the world, the better chance of being able to identify that individual based on gaze patterns. Second, two of the prior studies examined identification of the same target individual (Keles et al., 2022; Rigas & Komogortsev, 2014), while the other examined identification of twin pairs (Kennedy et al., 2017). It may be expected that accuracy for identifying the same target individual would be higher than identifying an individual’s twin. However, for the prior studies that examined identification of the same target individual (Keles et al., 2022; Rigas & Komogortsev, 2014), comparing accuracy between studies may be limited since those prior studies utilized a small number of dynamic movie stimuli. In contrast, the current work examines a large range of static pictures of complex scenes. An open question for future work should be examining the utility of gaze fingerprinting for dynamic movie stimuli versus static pictures. While the prior studies showcase some precedence for studying the general concept of gaze fingerprinting, the current study is the first to examine it in a large number of participants and at large-scale with respect to the range of stimuli. The proof-of-concept represented in this study is that gaze fingerprinting at very large scales likely has very high potential as a personalized marker with high biological and translational significance. Furthermore, with the emergence of rapidly developing technologies that utilize virtual reality and remote eye tracking at very large scales, the potential for gaze fingerprinting will be maximized. Such a situation may open up a range of issues regarding exploitation of such personalized inference from big data and also data protection and security issues.

When compared to gaze fingerprinting employed at the level of single stimuli, this study shows some added utility for averaging gaze similarity for fingerprinting over a large range of stimuli. However, averaging procedures may also risk potentially hiding unique and important information for the purpose of gaze fingerprinting. This point has been underscored in the past with analogous literature in functional neuroimaging that contrasts mass univariate analysis to multivariate pattern decoding techniques (Haxby et al., 2001). Analogous to the stimulus-rich experimental design strategies of representational similarity analysis (RSA) (Kriegeskorte et al., 2008), the stimulus-rich aspect of our gaze fingerprinting design allows us to fingerprint individuals on each stimulus. While accuracy rates for any one stimulus are generally much lower than averaging across stimuli, the gaze fingerprinting output per subject and stimuli represent unique ‘gaze fingerprint barcodes’ that when viewed as a pattern, rather than in isolation, may have important utility in the context of identifying individuals. Here we find that these ‘gaze fingerprint barcodes’ are sparse but are far from random.

We replicably find that around 94-95% of all individuals have a fingerprintable stimuli count that is significantly higher than the null distribution for fingerprintable stimuli count when subject identifier is randomly permuted. Clustering these gaze fingerprint barcode patterns shows that each individual is its own unique cluster. Thus, our findings confirm that these non-random sparse codes are indeed an identifiable signature per individual when taken into account as a pattern across a stimulus-rich design. Further bolstering the idea that gaze fingerprint barcodes are stable non-random individuating signatures, we showed in follow-up pre-registered experiments that barcodes are stable over long periods of time, from weeks to months to years and that barcode similarity does not heavily vary as a function of time. It is notable that such barcoding output is not possible without a stimulus-rich sampling of gaze across a large array of visual stimuli. However, the 700 stimuli utilized here is but a fraction of the total stimuli that possibly reveal unique individuating signatures for how individuals uniquely view the world. Thus, there is likely vast potential for expanding on this approach with much larger-scale datasets and expansion into dynamic stimuli (e.g., naturalistic movie viewing). There is also likely large potential for taking a ‘thin slices’ approach (Byrge & Kennedy, 2019) to identifying what would be the smallest amount of data across stimuli that would be useful for gaze fingerprinting. Such a ‘thin slice’ approach is beyond the scope of the current work but would be important to explore in future research.

Finally, we have shown that gaze fingerprint metrics are not only relevant for identifying individuals but can also be used to predict variance in other important phenotypic traits of relevance to conditions like autism. Atypical gaze patterns have long been studied in autism (Constantino et al., 2017; Jones & Klin, 2013; Wang et al., 2015) and may represent a key example of how genetic starting points linked to autism affect brain development in ways to create atypical niche construction via how individuals sample the world around them from very early points in development (Johnson, 2017; Johnson et al., 2015; Klin et al., 2020; Lombardo et al., 2019; Shultz et al., 2018). Both individual-sensitive gaze patterns and autistic traits are likely phenotypes sensitive to heritable polygenic or omnigenic common genetic architecture (Constantino et al., 2017; Constantino & Todd, 2003; Kennedy et al., 2017; Portugal et al., 2023; Robinson et al., 2016) and support the idea that gaze fingerprinting metrics may be a genetically sensitive signal that can predict variation in autistic traits. Here we find evidence supporting the utility of gaze fingerprinting metrics as predictive of variation in autistic traits such that higher gaze fingerprintability is linked to lower levels of autistic traits. This finding matches up to some evidence supporting that gaze patterns are more idiosyncratic in autism and linked to phenotypic variability in symptom severity (Avni et al., 2020). Within autism research, some have considered the hypothesis of enhanced neural and/or behavioral noise as a key feature of the autistic brain that leads to idiosyncrasy (Avni et al., 2020; Benkarim et al., 2021; Bleimeister et al., 2024; Bolton et al., 2018; Byrge et al., 2015; Cavallo et al., 2018; Dinstein et al., 2012, 2015; Hahamy et al., 2015; Hasson et al., 2009; Keles et al., 2022; Milne, 2011; Nunes et al., 2019; Pegado et al., 2020; Rubenstein & Merzenich, 2003; Sohal & Rubenstein, 2019; Trakoshis et al., 2020). Future work could examine whether enhanced neural noise leads to more idiosyncratic or less consistent gaze patterns within an individual that then related to lower gaze fingerprintability.

In summary, we show that gaze fingerprinting is possible both by averaging gaze similarity across stimuli, but also by preserving the gaze fingerprint information held within gaze fingerprint barcodes. The insights of this study were enabled by the use of a stimulus-rich design and are supported by replicable evidence across discovery and replication sets as well as confirmation of the non-random nature of gaze fingerprint barcodes from pre-registered longitudinal follow-up experiments over short and long time frames. Such work is important for future work decomposing individually-sensitive information within gaze patterns and may aid translational work related to understanding atypical gaze patterns in conditions like autism.

## Supporting information

Suppmentary Tables

## Research Transparency Statement

### General Disclosures

#### Artificial intelligence

No artificial intelligence assisted technologies were used in this research or the creation of this article.

#### Ethics

This study was approved by the local ethics committee (Comitato Etico per le sperimentazioni cliniche dell’Azienda Provinciale per Servizi Sanitari della Provincia Autonoma di Trento; APSS). All participants gave written informed consent and were reimbursed for their time participating in the study.

---

**Supplementary Figure 1:**
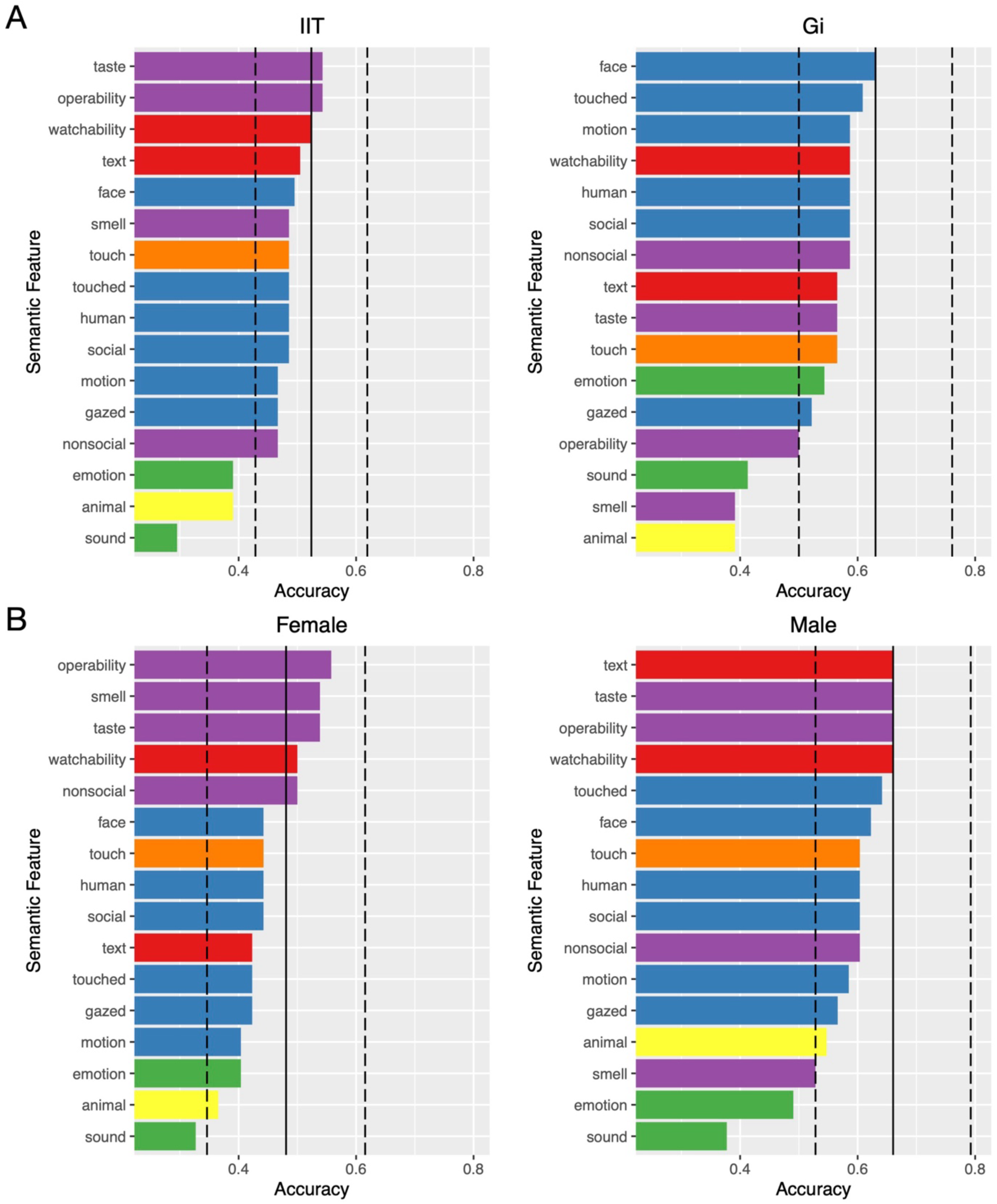
Analysis of IIT and Gi samples and then on IIT sex-stratified samples. Panel A shows results of gaze fingerprint accuracy (x-axis) on IIT (left) and Gi (right) samples. Panel B shows sex-stratified analyses of the IIT sample (females, left; males, right). The vertical black lines in each panel depict gaze fingerprint accuracy when average gaze similarity is computed across all 700 stimuli (dotted vertical black lines represent bootstrap 95% confidence intervals). Each bar shows gaze fingerprint accuracy when computed as average gaze similarity on subsets of stimuli grouped by semantic features shown on the y-axis.

**Supplementary Figure 2:**
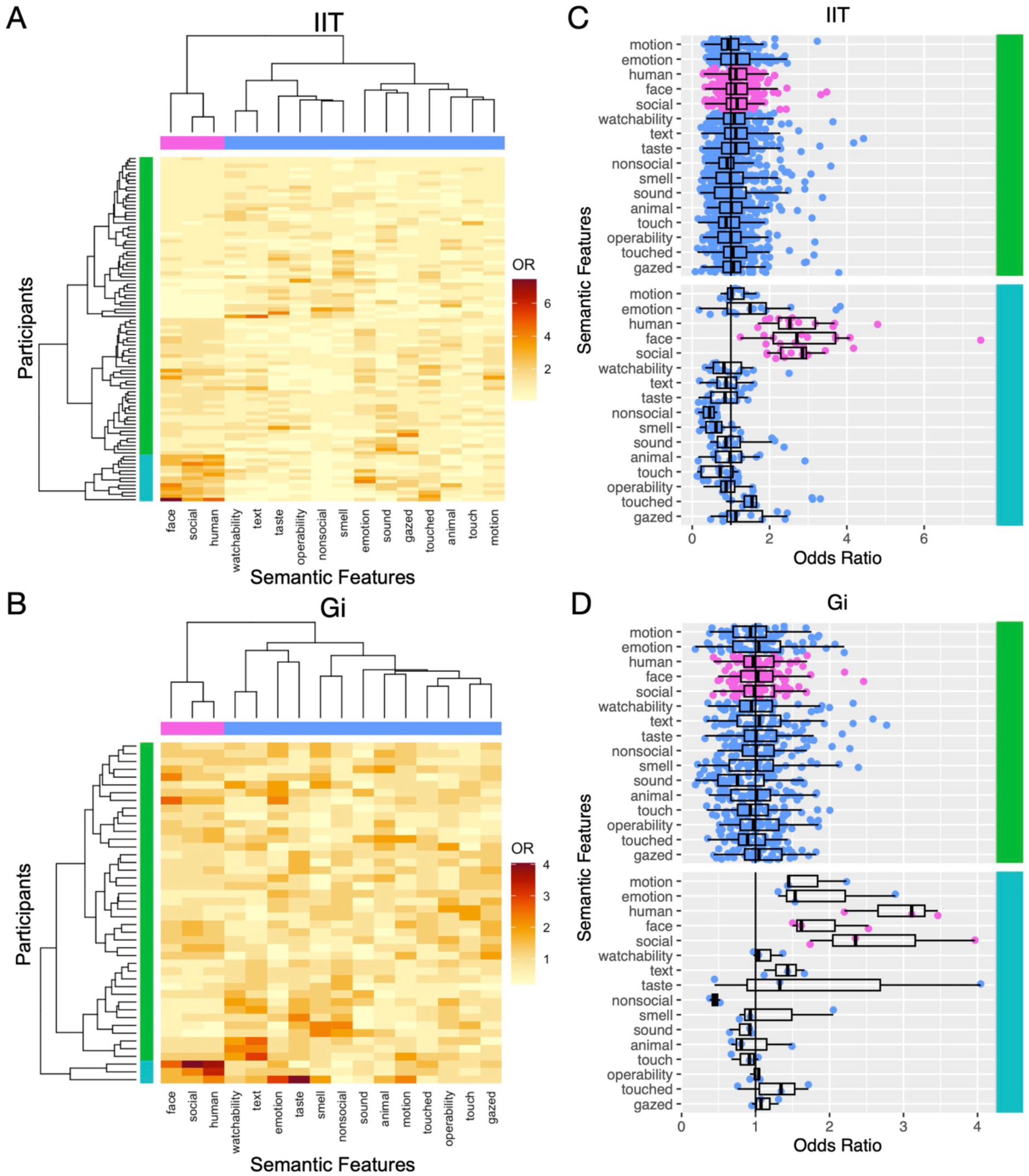
Semantic feature representation within gaze fingerprint barcodes signatures. Panels A (IIT discovery) and B (Gi replication) show heatmaps whereby color represents semantic feature enrichment odds ratios (OR) computed from fingerprintable stimuli sets of each subject. For each subject (rows), we computed enrichment odds ratios, which reflect an effect size relevant to the degree of overlap between a subject’s fingerprintable stimuli and stimuli of a specific semantic feature category (columns). Higher enrichment ORs indicate greater overlap between a subject’s fingerprintable stimuli and stimuli from a particular semantic feature. Rows of these enrichment OR matrices represent individual subject’s ‘semantic feature enrichment barcode’ and are clustered to identify subtypes with similar patterns of semantic feature enrichment. Columns of this matrix are also clustered to identify collections semantic features with similar enrichment profiles across subjects. Panels A and B show that the turquoise cluster along the subject dimension (rows) can be replicably identified with hotspots that tend to show higher enrichments for human, face, and social features. Panels C (IIT discovery) and D (Gi replication) show odds ratios (x-axis) for each feature (y-axis) and where each individual is a dot. The top and bottom panel of these plots represent the turquoise and green subtypes indicated in panels A and B.

